# Integrative network modeling of colorectal cancer reveals diagnostic signatures and therapeutic targets

**DOI:** 10.1101/2025.08.18.670822

**Authors:** Faiz M. Khan, Muhammad Naveez, Ali Salehzadeh-Yazdi, Vineetha Rajendran, Olaf Wolkenhauer, Julio Vera Gonzalez

## Abstract

Emerging evidence suggests that the interplay between multiple signaling pathways and the immune microenvironment influences tumorigenesis in cancers such as colorectal cancer (CRC). To gain an in-depth understanding of CRC mechanisms and identify novel therapeutic targets, we constructed a molecular interaction map (MIM) integrating key signaling pathways from cancer cells and immune cells within the tumor microenvironment. This map comprises 218 molecules and 328 interactions, curated using PubMed references and official gene symbols. We dynamically simulated the MIM for individual pathways and combinations via stimulus-response and perturbation analyses, calibrating the model with two CRC datasets: GSE1323 (primary tumor-to-metastasis progression) and GSE8671 (normal mucosa-to-adenoma progression), which served as experimental conditions. Simulations revealed distinct disease signatures for GSE1323, including (1) simultaneous activation of TNF/TNFRSF1A,B and EGF/EGFR with inactivation of ERE/ESR, and (2) simultaneous activation of TNF/TNFRSF1A,B and TLR4 with inactivation of ERE/ESR. For GSE8671, the signature was simultaneous activation of TNF/TNFRSF1A,B and TLR4. In silico perturbation analysis identified potent anti-cancer effects from concurrent inhibition of MAPK3 and STAT3 (GSE1323), and ELK1/ATF2 and STAT3 or MAPK14 and STAT3 (GSE8671), significantly reducing epithelial-mesenchymal transition (EMT), proliferation, and inflammation while increasing apoptosis. Disease signatures and therapeutic targets were validated using patient data through Kaplan-Meier survival analysis and machine learning. This integrative model recapitulates cancer biology, predicts biomarkers and therapeutic targets, and is extensible to other immunogenic cancers.

## Introduction

Cancer progression involves various (epi)genetic malfunctions that result in the dysregulation of single or multiple cellular pathways governing essential processes such as proliferation, cell death, inflammation, and migration [1]. Inflammatory and cancer-related pathways including tumor necrosis factor (TNF) [2], transforming growth factor beta (TGFB) [3], toll-like receptor 4 (TLR4) [4], Wingless/Integrated (WNT) [5], epidermal growth factor (EGF) [6], and estrogen signaling [7]; have all been shown to contribute to colorectal cancer (CRC) progression. These pathways are interconnected and influence each other, producing coordinated regulatory effects on cellular functions. While significant progress has been made in understanding the roles of individual pathways in CRC, the combined effects of multiple pathways on disease progression remain insufficiently explored.

To address this, we constructed, curated, and annotated a molecular interaction map (MIM) that integrates TNF, TGFB, TLR4, WNT, EGF, and estrogen pathways, revealing extensive cross-talk among them. To further investigate the interplay between CRC cells and immunity, we incorporated the inflammatory microenvironment network from Lu et al. [8] into our map. The resulting MIM contains state-of-the-art knowledge about 328 causal interactions among 218 molecules, detailing their roles in regulatory processes such as inflammation, epithelial-mesenchymal transition (EMT), proliferation, and apoptosis. The map is publicly available at https://zenodo.org/record/7104858.

Large-scale MIMs (disease maps) have become popular for computationally encoding molecular information about genes, proteins, and pathways involved in disease [9–14]. MIMs are machine-readable, facilitating standardization, exchange, and computational analysis to identify key regulatory players and potential therapeutic targets. For example, our previous work on the E2F1 MIM [9] in cancer successfully identified molecular mechanisms of EMT in bladder and breast cancer, validated through patient data and in vitro experiments. Other notable MIMs include the Parkinson’s disease map [12] and the Atlas of Inflammation-Resolution (AIR) [10].

MIMs also provide a scaffold for the dynamic analysis of complex biological systems [9]. Regulatory relationships among nodes can be translated into mathematical equations to describe temporal changes in molecule concentrations or expression levels. Systems biology models range from qualitative logic-based models (e.g., Boolean models) to quantitative models using ordinary differential equations [15]. Boolean models, which represent each node as either active (1) or inactive (0) [16,17], are computationally simple and do not require detailed quantitative data, yet can reveal mechanisms driving complex system behaviors [18–20]. They have been applied to explain processes such as the cell cycle [21–23], cell differentiation [24,25], cell death [26–28], and migration, and to identify pathway alterations [29,30] and combinatorial drug targets using in silico perturbations [31–33]. Patient-specific data can be integrated to create personalized models that reproduce disease phenotypes and predict therapeutic responses [34]. Previous studies have demonstrated the value of Boolean and logical modeling in colorectal cancer research. For instance, Nagaraj & Reverter (2011) [35] introduced a Boolean-based systems biology framework that integrated genetic, transcriptomic, and molecular data to identify novel cancer-associated genes and candidate biomarkers in CRC, showcasing the utility of Boolean logic for gene prioritization and network analysis. Lu et al. (2015) [8] constructed a Boolean network to explore inflammation-driven mechanisms in colitis-associated colon cancer, using attractor analysis and in silico perturbations to reveal regulatory modules and potential combinatorial drug targets, with some predictions validated experimentally. Recently, Béal et al. (2021) [36] developed personalized logical models to predict cancer response to BRAF inhibition in both melanoma and CRC, demonstrating the potential of logic-based approaches for individualized therapy prediction by integrating cell line-specific omics data and simulating drug responses.

In this study, we encoded our CRC map into a Boolean model to dynamically analyze the effects of individual and combined pathways using stimulus-response and perturbation analyses. Boolean functions were assigned to each node, determining its state based on its regulators via logical gates (AND, OR, NOT), and calibrated with fold-change expression data from two CRC datasets: GSE1323 [37] and GSE8671 [38,39]. For each dataset, an individual Boolean model was constructed, comprising three layers: (1) input (ligand-receptor nodes), (2) regulatory (signaling cascades and transcriptional events), and (3) phenotype (cell fate outputs). The models are available at https://zenodo.org/record/7143340#.Yzx2L4RBwuU.

The input layer was initialized using fold-change values from the respective datasets, and logical steady states were calculated for each node by propagating signals through the network using CellNetAnalyzer (CNA) [40]. This approach enabled us to obtain stimulus-response patterns and perform systematic perturbations, revealing how signals propagate and ultimately affect phenotypes such as proliferation, apoptosis, invasion, and inflammation. Consistent with the literature, CRC is characterized by sustained proliferation, migration, inflammation, and limited apoptosis [1,41–43]. For GSE1323, stimulus-response simulations identified two regulatory signatures linked to high inflammation, invasion, proliferation, and low apoptosis: (1) simultaneous activation of TNF/TNFRSF1A,B and EGF/EGFR, with inactivation of ERE/ESR; and (2) simultaneous activation of TNF/TNFRSF1A,B and LY96/TLR4, with inactivation of ERE/ESR. For GSE8671, simultaneous activation of TNF/TNFRSF1A,B and LY96/TLR4 led to similar phenotypic outcomes.

Perturbation analysis identified critical nodes whose inhibition (gain-or loss-of-function) significantly influenced disease phenotypes, highlighting potential diagnostic and therapeutic targets. For GSE1323, inhibition of MAPK3 and STAT3 profoundly reversed malignant phenotypes, while for GSE8671, double inhibition of ELK1/ATF2 and STAT3 or MAPK14 and STAT3 had similar effects. These in silico predictions are consistent with experimental observations [2,6,44–52]

To further validate our findings, we performed Kaplan-Meier survival analysis using TCGA CRC data and demonstrated that patients can be stratified into different survival groups based on our predicted signatures (Figure 2). Additionally, supervised machine learning models (k-nearest neighbor, support vector machine, naïve Bayes, and random forest) trained on gene expression data for the predicted signatures effectively distinguished healthy from tumor samples, supporting their predictive utility (Figure 4).

**Figure 1.**
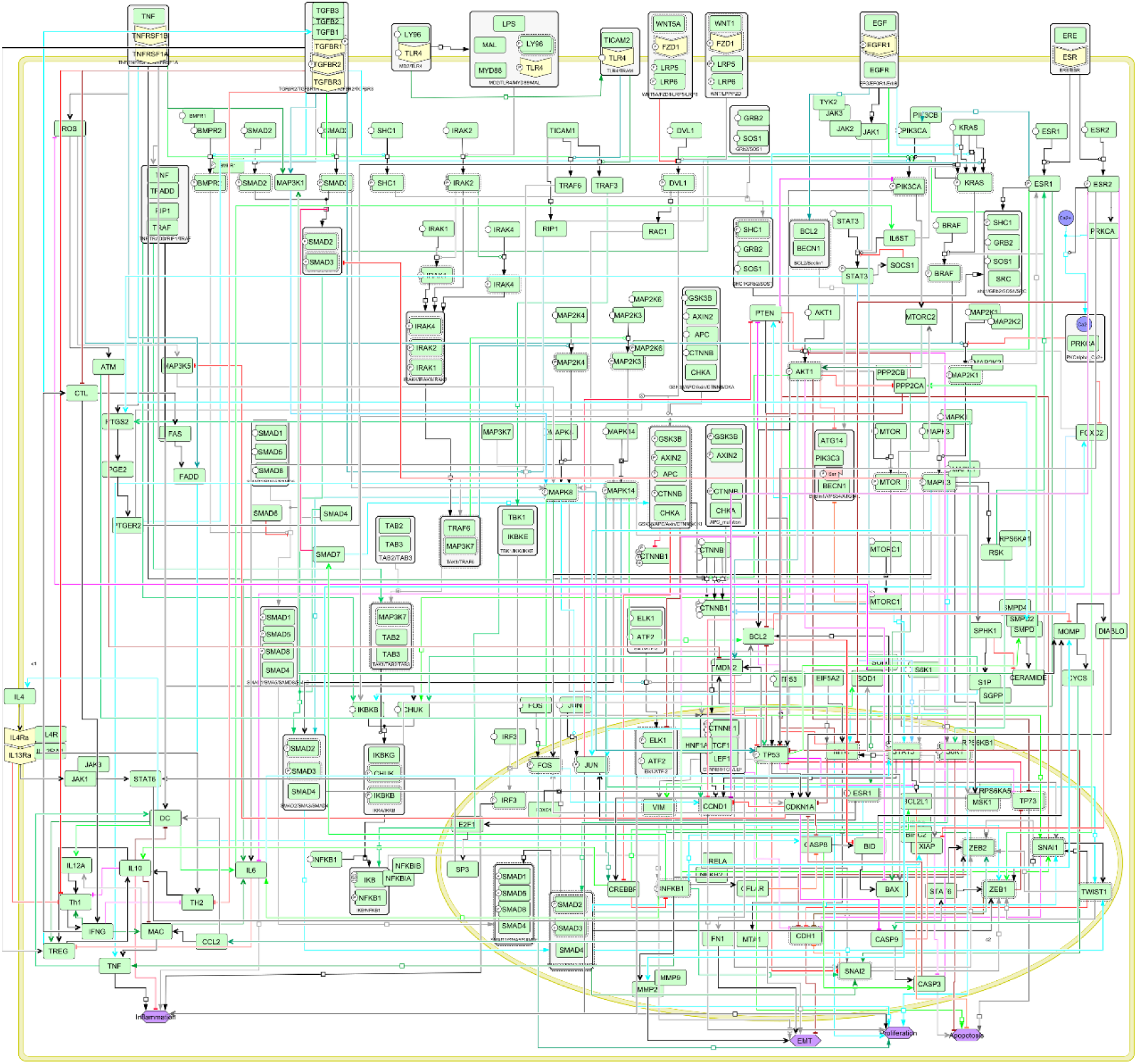
CellDesigner visualization of the colorectal cancer (CRC) progression map. Reddish edges (red, pink, magenta) indicate inhibitory regulatory interactions, while black, blue, cyan, and gray edges denote activation. The map comprises 218 nodes and 328 edges, with nodes and edges annotated by official names and PubMed IDs, respectively. The complete map is available for download at https://zenodo.org/record/7104858.

**Figure 2.**
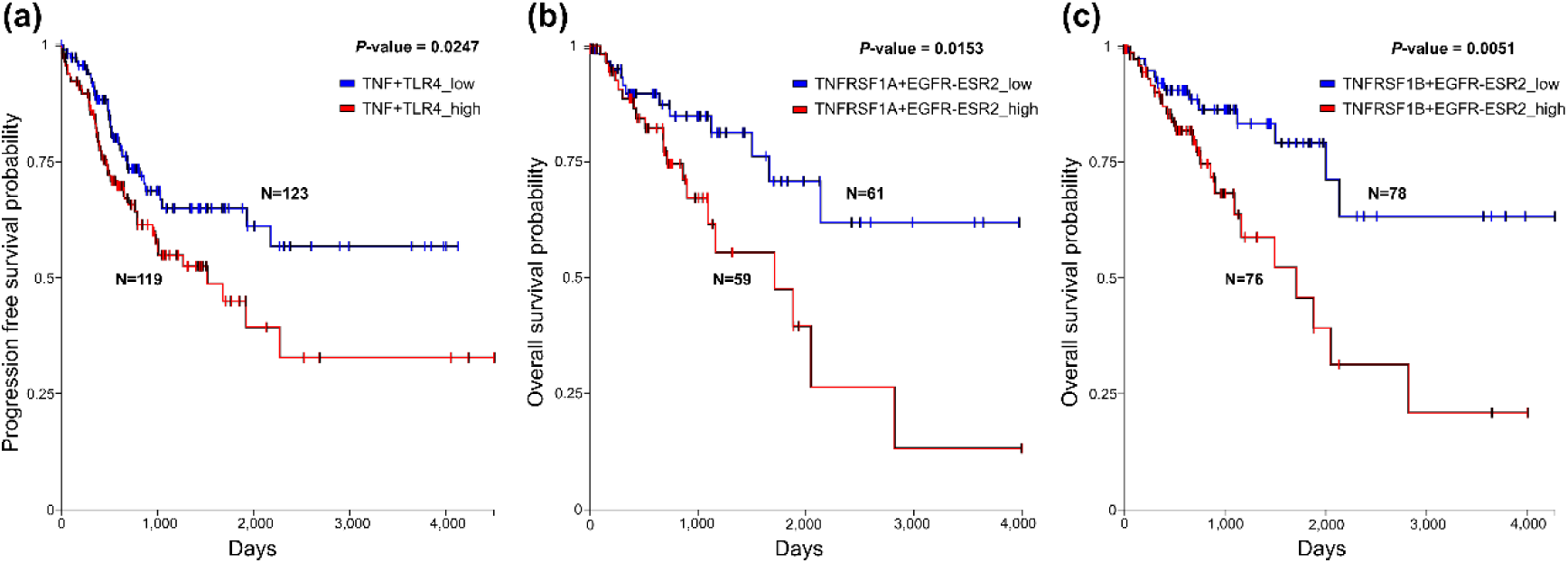
Validation of in silico predicted disease signatures using TCGA data and Kaplan-Meier survival analysis. Survival plots demonstrate that patients with high expression of the predicted signatures exhibit significantly lower survival probabilities compared to those with low expression, supporting the prognostic value of the model-derived signatures [83].

**Figure 3.**
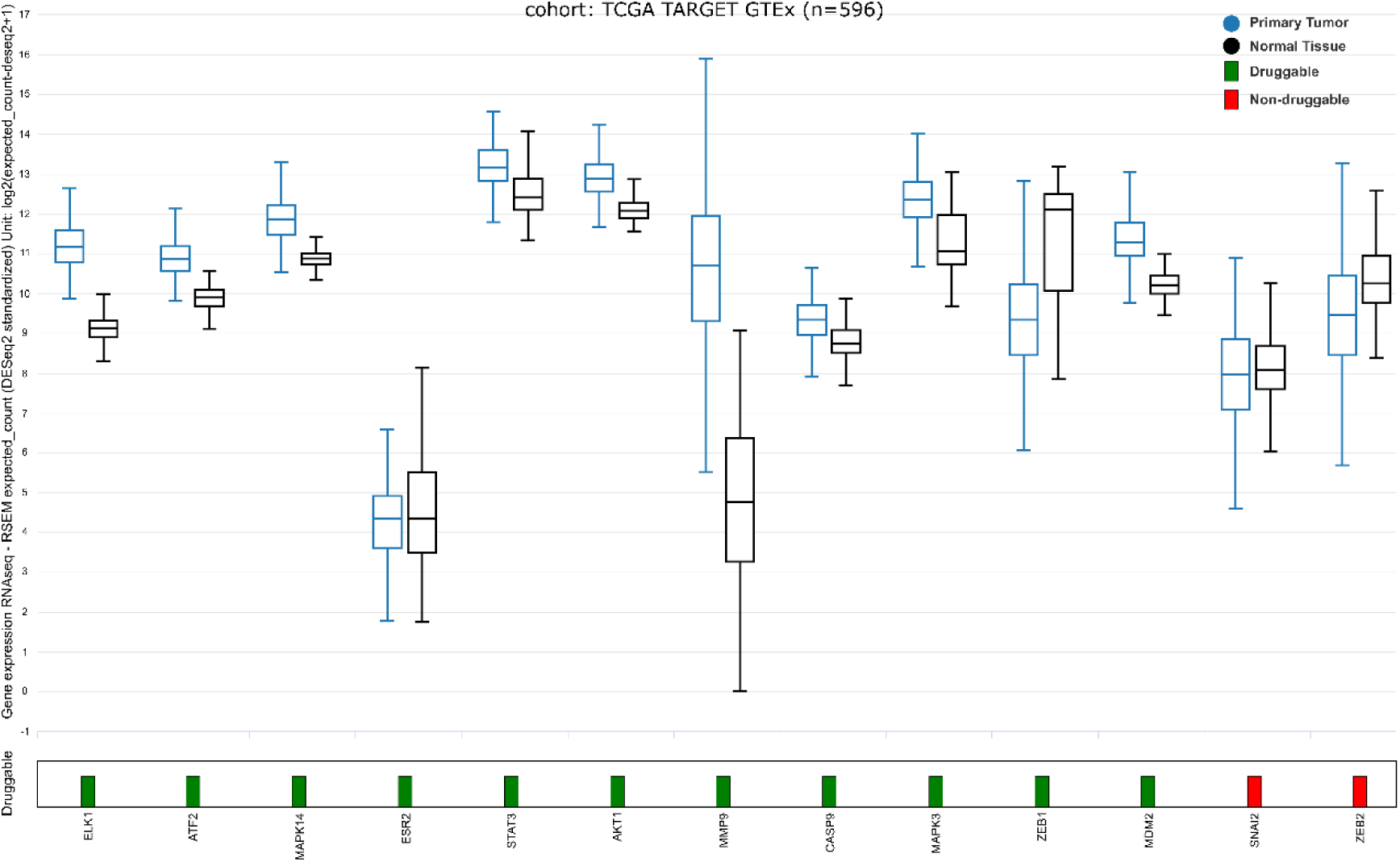
Comparison of predicted drug target expression in primary tumor versus normal tissue using TCGA and GTEx data (source: https://xenabrowser.net/). Box plots show that genes identified as combination drug targets generally have higher expression in primary tumors (blue boxes) compared to normal colon tissue (black boxes), while ESR2 displays lower expression in tumors, consistent with model predictions. The bar chart at the bottom summarizes drug availability for each predicted target in colorectal cancer: green bars indicate available drugs, and red bars indicate no available drugs (source: https://www.genecards.org/).

**Figure 4.**
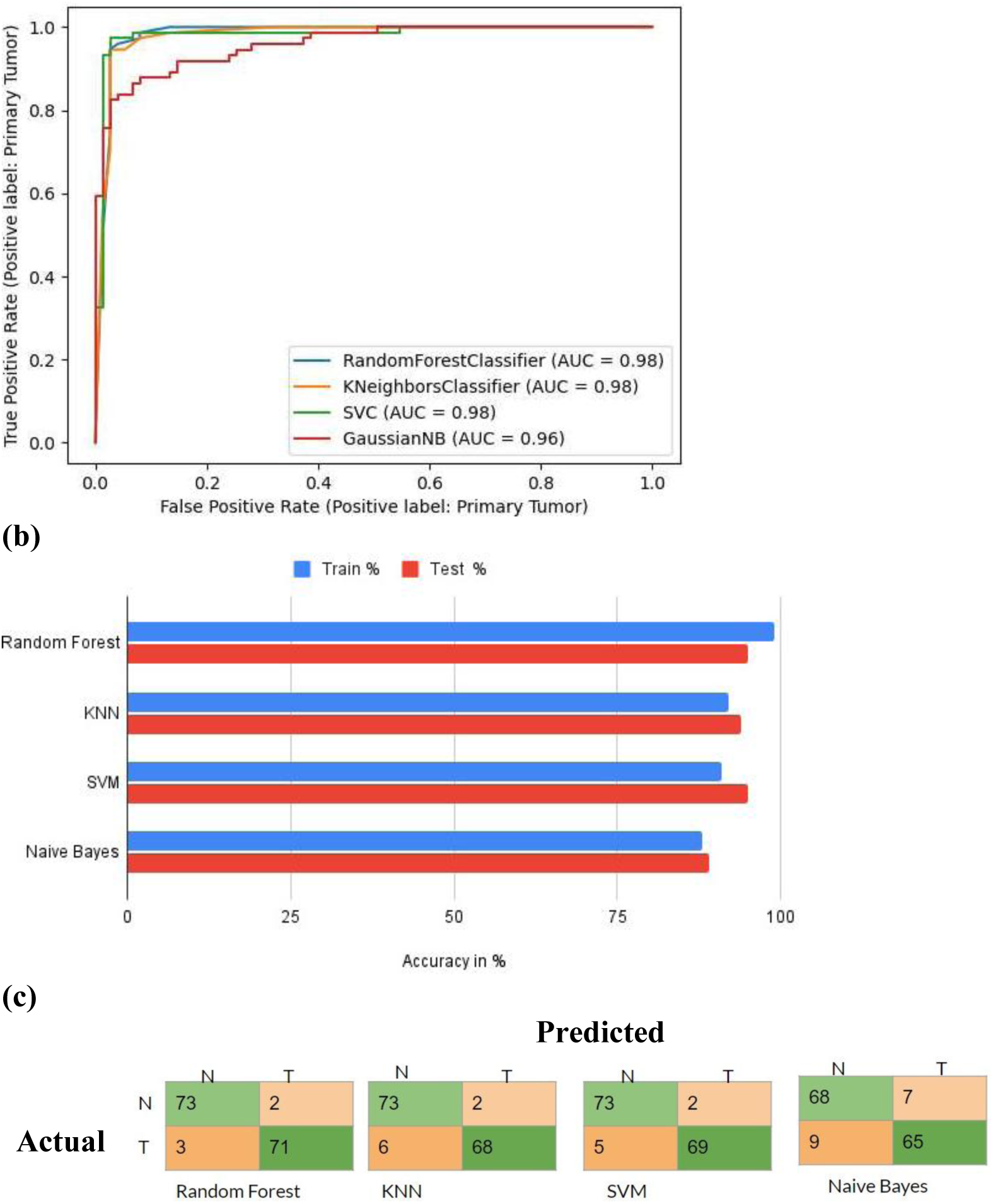
(a) Receiver operating characteristic (ROC) curves for binary classification of the training dataset using four classifiers: support vector machine (SVC), k-nearest neighbor, Naïve Bayes (GaussianNB), and random forest. **(b)** Comparison of classifier accuracy for both training (blue) and test (red) datasets. **(c)** Confusion matrices for the different classifiers, based on cross-validation results with the test dataset. N: normal healthy tissue; T: tumor tissue.

In summary, our study provides an integrated view of multiple signaling pathways and the inflammatory microenvironment in CRC progression. The map and model presented here offer a valuable resource for mechanistic studies, disease signature identification, and the development of novel combinatorial therapies

## Results

### The MIM represents multiple pathways regulating CRC progression

We manually constructed and curated a molecular interaction map (MIM) representing the regulatory events underlying colorectal cancer (CRC) progression by retrieving pathway information from extensive literature searches (Figure 1). The map encompasses the TNF, TGFB, TLR4, WNT, EGF, and Estrogen signaling pathways, all of which are recognized as central drivers of CRC pathogenesis and progression [3,53,54]. These pathways regulate key cancer phenotypes, including proliferation, apoptosis, inflammation, and migration, and are known to interact extensively, forming a highly interconnected signaling network [3,53,55].

The MIM captures events ranging from ligand-receptor binding and activation of downstream signaling cascades to the induction of relevant transcription factors and their gene targets. To further incorporate the influence of the inflammatory tumor microenvironment, we integrated the immune network developed by Lu et al., providing a more comprehensive representation of CRC biology.

The final map contains 218 nodes representing biological entities such as genes, proteins, complexes, or transcription factors and 328 edges, which denote regulatory relationships including phosphorylation, ubiquitination, activation, inhibition, or transcriptional activation. All nodes and edges are annotated with official gene/protein names and corresponding PubMed references, ensuring traceability and reproducibility. The MIM is publicly accessible at https://vcells.net/colorectal-cancer/, and the CellDesigner file can be downloaded from https://zenodo.org/record/7104858.

A brief description of each pathway is provided in the Supplementary Material. This resource lays the groundwork for dynamic modeling and simulation of CRC signaling, enabling in-depth analysis of individual and combined pathway effects and facilitating the identification of diagnostic and therapeutic targets [55].

### Logic-based model construction of CRC

To systematically identify disease signatures and therapeutic targets in colorectal cancer (CRC), we encoded the molecular interaction map (MIM) as a logic-based (Boolean) model. This approach enables simulation-based stimulus-response and perturbation analyses, allowing us to explore the dynamic behavior of CRC signaling networks under various conditions. The model comprises 452 Boolean functions, each describing the regulatory relationship (activation or inhibition) between upstream regulators and downstream targets using logical operators (AND, OR, NOT; see Figure 5 and Materials and Methods).

**Figure 5.**
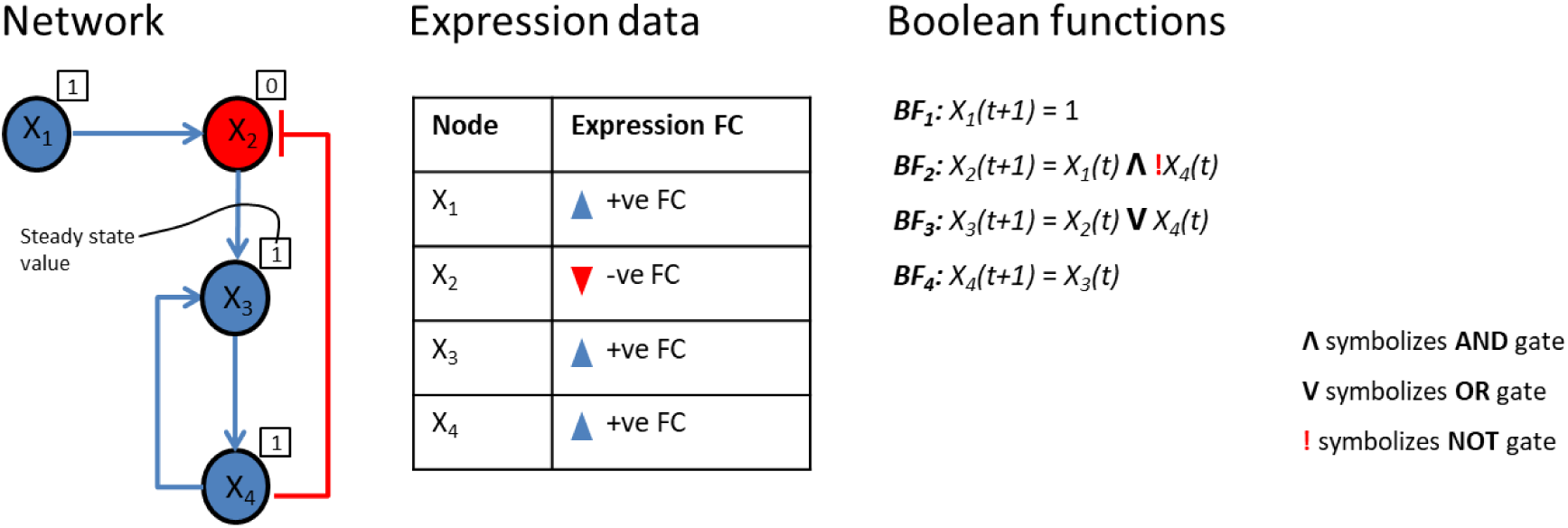
Construction and calibration of a Boolean toy model. The network consists of four nodes (X_1_, X_2_, X_3_, and X_4_), with regulatory relationships depicted by arrows for activation and T-shaped lines for inhibition. Node X_1_, lacking upstream regulators and displaying a positive fold-change (FC) value, is initialized to state ‘1’ (see BF_1_). Node X_2_ is regulated by X_1_ (activator) and X_4_ (inhibitor); although both regulators have positive FC values, X_2_’s negative FC indicates a collective regulatory effect, encoded with an AND gate (see BF_2_). Node X_3_ is regulated by X_2_ and X_4_ (both activators); with X_2_ showing a negative FC and X_4_ a positive FC, and X_3_ itself having a positive FC, this independent regulatory effect is represented by an OR gate (see BF_3_). Node X_4_ is activated solely by X_3_, and both have positive FC values, so X_4_’s future state depends on X_3_’s present state (see BF_4_). After initializing the input node, simulation ensures that the steady-state value of each node (shown as a box over the node with 0 or 1) matches its FC expression value (red for negative, blue for positive); thus, nodes with positive FC (blue) correspond to a steady-state value of 1, and those with negative FC (red) correspond to 0.

To facilitate analysis, the model was structured into three layers: (1) the input layer, containing ligand-receptor nodes (e.g., TNF/TNFRSF1A,B; TGFB1,2/TGFBR1,2,3; TLR4/LY96; WNT1,5/FZD1/LRP5,6; EGF/EGFR1/EGFR; and ERE/ESR) that mimic cell-environment interactions; (2) the regulatory layer, encompassing the signaling cascades and transcriptional events triggered by ligand-receptor interactions; and (3) the output layer, representing key CRC phenotypes such as inflammation, apoptosis, epithelial-mesenchymal transition (EMT), and cell proliferation. The steady-state value of each phenotype node reflects the long-term cellular response to specific stimuli or perturbations.

The Boolean models, provided in CNA format at https://zenodo.org/record/7143340#.Yzx2L4RBwuU, were calibrated using fold-change (FC) expression data from two publicly available CRC datasets: GSE1323 [37] and GSE8671 [38]. Model calibration involved manually matching the FC expression values of source and target nodes to the regulatory relationships defined by their respective Boolean functions (see Figure 5).

For simulation, we initialized the input layer by assigning states to ligand-receptor nodes based on the FC expression values: a node was set to’1’ if any constituent gene had a positive FC, and’0’ otherwise. Due to data sparsity, this conservative approach ensured robust initialization. We then iteratively refined the Boolean functions governing node transitions, retaining those that accurately reproduced the observed FC data in the regulatory layer during simulations. In this framework, negative FCs correspond to state’0’ and positive FCs to state’1’ [9,18].

This logic-based modeling strategy enables the integration of complex pathway crosstalk and provides a computational platform for predicting the effects of targeted perturbations on CRC phenotypes, in line with recent advances in systems biology and cancer modelling [56–58].

### Stimulus-response analysis of the model

After model calibration, we simulated the model for all possible combinations (around 128) of ligand-receptor nodes and calculated the steady-state value for each node in the model. This allowed us to identify the effect of each ligand-receptor node, both individually and in combination, on the phenotypes (Table 1a and 1b). Under the experimental condition of the dataset GSE1323, our simulations suggest two ligand-receptor signatures associated with high inflammation, invasion, proliferation, and low apoptosis in CRC progression: (1) simultaneous activation of TNF/TNFRSF1A,B and EGF/EGFR1/EGFR with inactivation of ERE/ESR ligand-receptor nodes; and (2) simultaneous activation of TNF/TNFRSF1A,B and LY96/TLR4 with inactivation of ERE/ESR ligand-receptor nodes (Table 1a). For the dataset GSE8671, we found that the simultaneous activation of TNF/TNFRSF1A,B and LY96/TLR4 can similarly lead to high inflammation, invasion, proliferation, and low apoptosis in CRC progression (Table 1b).

**Table 1.**
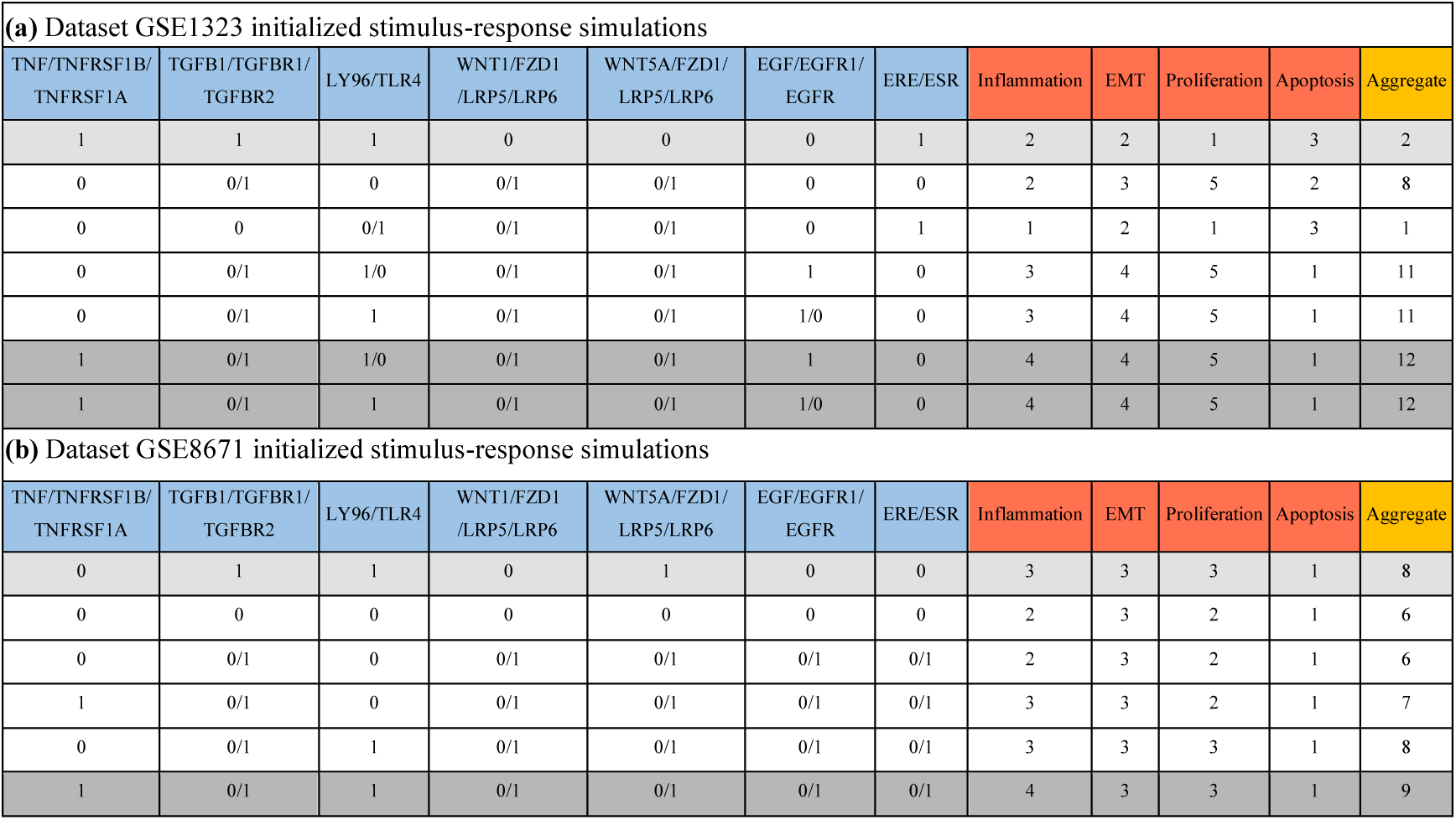
Summary of stimulus-response simulation results for models trained with datasets (a) GSE1323 and (b) GSE8671. Ligand-receptor nodes (blue) are encoded as Boolean (0 or 1) and represent input conditions; their effects on phenotypes (red) - inflammation, epithelial-mesenchymal transition (EMT), proliferation, and apoptosis - are modeled with multi-valued logic (0-5), reflecting the number of active regulators. The aggregate (yellow) is calculated by subtracting the steady-state value of apoptosis from the sum of inflammation, EMT, and proliferation, providing a composite measure of overall effect. For nodes labeled 0/1, simulation results are the same whether the node is 0 or 1. The light gray row indicates the initial condition based on the dataset’s fold-change expression profile; dark gray rows at the bottom represent the combinations yielding the highest aggregate phenotype values.

In all identified signatures, TNF/TNFRSF1A,B is involved, suggesting that activation of the TNF pathway is a crucial factor for adverse CRC progression. This observation aligns with emerging experimental evidence showing that TNF pathway activation increases the proliferative and invasive potential of CRC, and that blocking TNF can inhibit cancerous phenotypes [2,45,46]. The CRC map (Figure 1) provides detailed molecular information, illustrating that TNF regulates multiple cellular processes within the immune microenvironment, cell survival, proliferation, and apoptosis. Specifically, TNF triggers cell survival and proliferation pathways including NFKB [2], PI3K/AKT [47], and JNK/p38/AP-1 [2,59]; as a pro-inflammatory cytokine, it activates immune response pathways involving cytotoxic T lymphocytes (CTL), COX2 [45,60], and dendritic cells (DC) [8,61,62]; and it inhibits apoptosis through the induction of XIAP, BCL2, and c-Flip [8,63].

Activation of the EGF/EGFR1/EGFR pathway in CRC is also supported by experimental observations [6,64]. Our map shows that EGFR activates multiple signaling pathways, including the RAS/RAF pathway, regulating proliferation and survival through MYC, RSK, MSK, SPHK1, and STAT3; the JAK/STAT pathway, activating MYC, CCND1, BCL2L1, SOD, IL6, SNAIs, VIM, and MMPs to regulate survival, anti-apoptosis, migration, and immune response; and the PI3K/AKT pathway, activating protein synthesis factor mTOR/S6K, immune response regulator NFKB, and apoptosis inhibitor BCL2.

Experimental evidence also suggests that activated TLR4 signaling is involved in many inflammation-driven malignancies, including CRC, and that inhibition of TLR4 ligands has been proposed as a promising anti-cancer therapeutic strategy [44,48]. Our map depicts the TLR4-regulated molecular machinery involved in immune responses and cell survival through NFKB, AP1, STATs, and IRFs signaling [44,48,65]. Furthermore, estrogen receptor beta (ESR2) has a protective role in CRC progression, and growing experimental evidence indicates that the loss of ESR2 is implicated in CRC progression [49–52].

### In silico perturbation analysis

To identify potential intervention targets capable of reversing deregulated phenotypes, we conducted in silico perturbation analyses for each disease signature. Our aim was to find nodes whose inhibition could suppress inflammation, epithelial-mesenchymal transition (EMT), and proliferation, while enhancing apoptosis. Perturbation experiments were performed by systematically changing the Boolean state (0 or 1) of each node in the regulatory layer. We explored both single and double perturbations to identify intervention targets that could reduce the aggregate phenotype to the minimum possible level.

For the dataset GSE1323, our simulation results suggest that simultaneous inhibition of MAPK3 and STAT3 can reduce the aggregate phenotype from 12 to 5. In the case of the dataset GSE8671, double perturbation of ELK1/ATF2 and STAT3 or MAPK14 and STAT3 can similarly reduce the aggregate phenotype from 9 to 4 (see Table 2a and b).

**Table 2.**
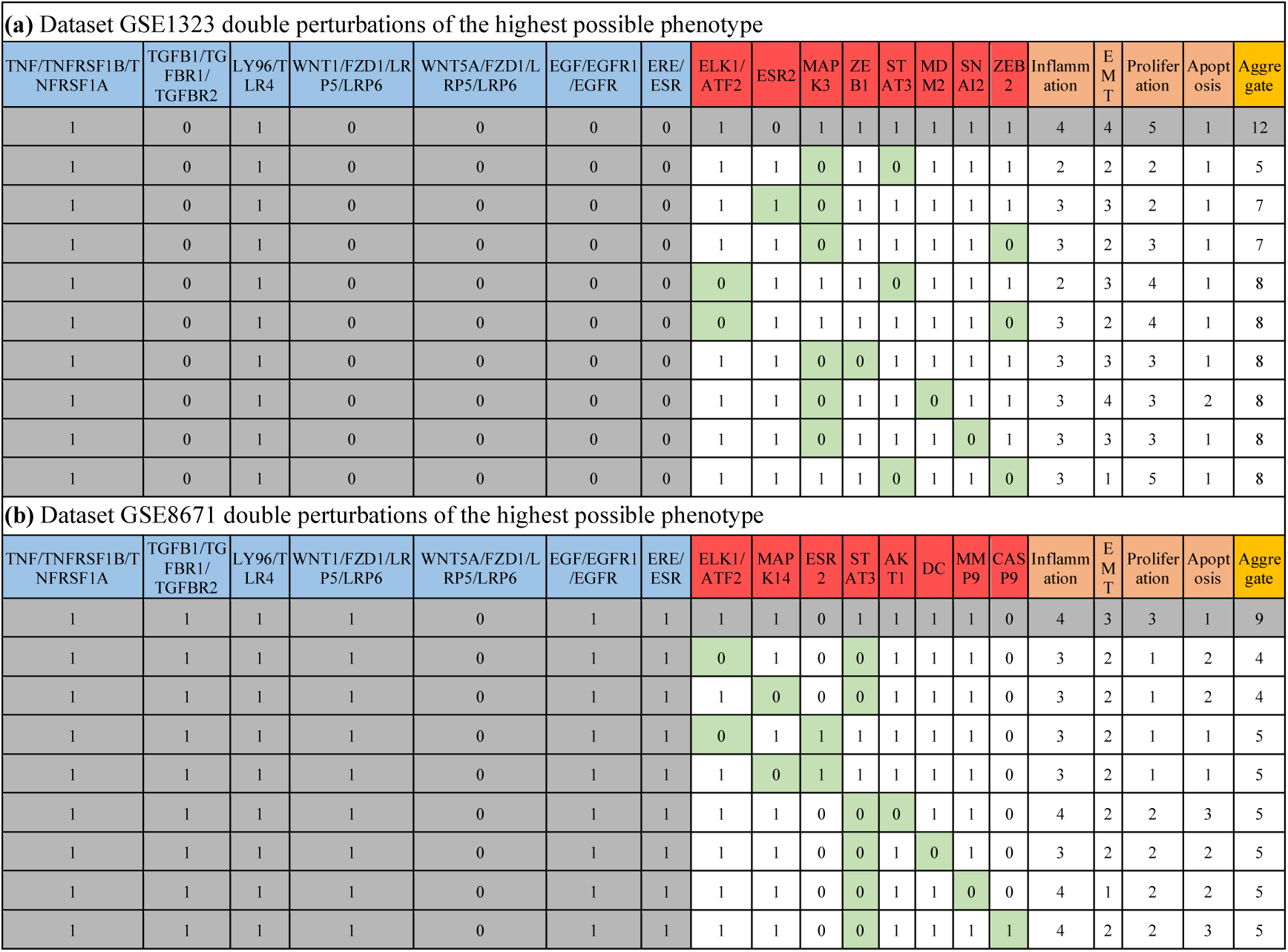
Double (two-node) perturbation of the highest possible phenotype for models trained with datasets **(a)** GSE1323 and **(b)** GSE8671. The highest possible phenotype and its corresponding disease signature, specifically, the ligand-receptor node configuration, are highlighted in dark gray. In the model simulations, the values for ligand-receptor nodes were kept fixed, while each node in the regulatory layer was systematically perturbed (from 0 to 1 or vice versa) to identify potential intervention markers. Green highlighted boxes in the table indicate the intervention markers that reduce the aggregate phenotype to the lowest possible level.

The in silico predicted drug targets are in agreement with previous experimental findings, where overexpression of STAT3 [66–69] and MAPK3 [70] has been associated with sustained proliferation and inflammation in CRC cells, and inhibition of these nodes could reduce proliferation and induce apoptosis [71–74]. Jin et al. [75] investigated a combination therapy of STAT3 inhibitor and MEK (the upstream regulator of MAPK3) inhibitor and found that the combination therapy has a stronger effect in suppressing cell viability and inducing apoptosis compared to monotherapy.

Similarly, previous experimental studies suggest that MAPK14 [76] and ELK1/ATF2 [77,78] can be critical markers for colon cancer and that their inhibition could be a potential therapeutic strategy [79–82].

### Validation of in silico predictions with patient data

To validate our in silico predictions, we utilized two colon cancer datasets from The Cancer Genome Atlas (TCGA): the TCGA Colon and Rectal Cancer (COADREAD) cohort (N=242 patients) (https://xenabrowser.net/datapages/?cohort=TCGA%20Colon%20and%20Rectal%20Cancer%20(COADREAD)&removeHub=https%3A%2F%2Fxena.treehouse.gi.ucsc.edu%3A443) and the TCGA Colon Cancer (COAD) cohort (N=120 patients) (https://xenabrowser.net/datapages/?cohort=TCGA%20Colon%20Cancer%20(COAD)&removeHub=https%3A%2F%2Fxena.treehouse.gi.ucsc.edu%3A443).

Kaplan-Meier survival analysis, performed using the UCSC Xena browser, stratified patients based on the expression profiles of our predicted disease signatures. For the GSE8671-derived signature (TNF/TNFRSF1A,B + LY96/TLR4) in the COADREAD cohort, patients were divided into low-expression (N=123) and high-expression (N=119) groups based on median TNF and TLR4 levels. This stratification revealed a significant survival difference (*p*-value=0.0247), with low-expression patients exhibiting higher progression-free survival probability than high-expression patients (Figure 2a).

For the GSE1323-derived signatures in the COAD cohort, we validated two patterns: First, for the signature involving TNFRSF1A/EGFR activation and ESR2 inactivation, stratification into low-expression (N=61) and high-expression (N=59) groups showed significantly higher overall survival (*p*-value=0.0153) in the low-expression group (Figure 2b). Second, for the signature combining TNFRSF1B/EGFR activation and ESR2 inactivation, low-expression patients (N=78) similarly demonstrated superior survival outcomes (*p*-value=0.0051) compared to high-expression counterparts (N=76) (Figure 2c).

These results confirm that our computationally predicted disease signatures effectively stratify colorectal cancer patients into clinically distinct survival groups, underscoring their biological and translational relevance. The validation across independent cohorts highlights the robustness of our integrative modeling approach, with survival curves generated using TCGA RNA-seq data and clinical annotations, group stratification based on median expression thresholds, and statistical significance assessed via log-rank tests. This establishes a foundation for future biomarker development and targeted therapeutic strategies in colorectal cancer

To further validate the intervention markers (drug targets) identified through our in silico perturbation analysis, we compared their expression profiles in primary tumor samples from TCGA cohorts with normal colon tissues from the GTEx database. Using the integrated’TCGA TARGET GTEx’ dataset accessed via the UCSC Xena browser (https://ucsc-xena.gitbook.io/project/how-do-i/tumor-vs-normal#using-the-tcga-target-gtex-study), we analyzed expression differences between tumor and normal tissues. The results demonstrated significantly higher expression of ELK1, ATF2, MAPK14, STAT3, AKT1, MMP9, MAPK3, MDM2, and SNAI2 in primary tumors, along with reduced expression of ESR2, consistent with our model predictions (Figure 3). While the expression patterns of CASP9 and ZEB1/2 did not fully align with our simulations, extensive experimental literature supports their relevance: low CASP9 expression correlates with CRC progression due to impaired apoptosis [84–86], and elevated ZEB1/2 expression drives epithelial-mesenchymal transition and metastasis [87–90]. Additionally, we assessed the druggability of these predicted targets using the GeneCards database (https://www.genecards.org/). This analysis confirmed that the majority of these markers, including STAT3, MAPK3, MAPK14, AKT1, and MDM2, are classified as druggable targets, with existing pharmacological inhibitors either in clinical use or under development (summarized in Figure 3, last row).

SNAI2 and ZEB2 are not directly druggable, but their upstream regulators offer promising indirect targets. SNAI2 expression is regulated by microRNAs including miR-200b-3p, miR-30c-1-3p, miR-363-3p, and notably miR-145, which suppress cancer invasiveness [91,92]. In colorectal cancer, inhibiting SNAI2 or restoring miR-145 enhances sensitivity to chemotherapy like 5-fluorouracil. Ubiquitin-related enzymes such as UbcH5c (UBE2D3) also modulate SNAI2 function epigenetically [93]. Meanwhile, ZEB2 is controlled by the miR-200 family (e.g., miR-200b, miR-200a, miR-429), which inhibit its transcription and EMT, offering indirect therapeutic avenues [94]. Targeting these regulatory networks may overcome metastasis and drug resistance in colorectal cancer. This druggability profile underscores the translational potential of our computational predictions for therapeutic development in colorectal cancer.

### Machine-learning based validation of disease signatures

In order to verify if the gene expression signatures obtained via network analysis and simulation has predictive power, we retrieved processed gene expression data from healthy tissue and tumor tissue samples and extracted the expression data for the investigated markers. We split the data into training (75%) and test (25%) datasets and tested several supervised ML models including random forest, KNN, SVM and Naïve Bayes. When considering the training data, the performance of the ML classifiers showed AU-ROC ranging from 0.96 to 0.98 (Figure 4a). When validating the performance of the models with the training data, all the ML models kept a high accuracy in the predictions (AU> 0.87, Figure 4b). The top-performing classifier for both training and validation data sets is a random forest. Also, the random forest model provided a more balanced accuracy when assessing the confusion matrices (Figure 4c).

## Discussion

Colorectal cancer (CRC) is characterized by the complex interplay of multiple deregulated genes and signaling pathways. In this study, we employed a network and logic-based modeling approach to systematically unravel the crosstalk between critical molecular factors involved in CRC progression. By manually constructing and curating a comprehensive molecular interaction map (MIM) that integrates major cancer signaling pathways and the immune microenvironment, we were able to simulate and analyze the dynamic behavior of CRC at a systems level.

The molecular interaction map (MIM) developed in this study is provided in standard SBGN format, with nodes annotated using official gene names. This ensures that the map is easily reusable and extensible to other pathways and diseases. Furthermore, the map supports visualization and integration of diverse data types, including mRNA, scRNA, and snRNA, enabling users to overlay experimental datasets and explore activity levels of key regulators and disease markers. An interactive version of the map is publicly available at https://vcells.net/colorectal-cancer/, where users can visualize expression data from datasets such as GSE1323 and GSE8671.

Our Boolean modeling and stimulus-response simulations identified distinct disease signatures that drive aggressive CRC phenotypes, including inflammation, proliferation, invasion, and resistance to apoptosis. Notably, signatures involving the simultaneous activation of TNF/TNFRSF1A,B and EGF/EGFR or TLR4, coupled with inactivation of ERE/ESR, were consistently associated with poor prognostic features. These findings are in agreement with experimental data, highlighting the central role of TNF, EGF, and TLR4 signaling, as well as the protective influence of ESR2, in CRC progression. However, the combinatorial effects of these markers warrant further experimental validation.

Perturbation analyses of our models revealed key intervention nodes, such as MAPK3, STAT3, ELK1/ATF2, and MAPK14, whose inhibition could robustly reverse malignant phenotypes in silico. Our proposed drug targets are supported by experimental observations; for example, many studies have investigated STAT3 combination therapies [75,95], and our analysis of patient-derived expression data showed elevated levels of these targets in primary tumors and confirmed their druggability. However, the efficacy of inhibiting STAT3 in combination with MAPK3, ELK1/ATF2, or MAPK14 remains to be experimentally validated and may represent a promising avenue for CRC therapy.

Extending previous CRC Boolean modeling efforts [8,35,36], our approach integrates a broader and more comprehensive set of CRC-relevant signaling pathways, systematically maps network dynamics to clinically meaningful phenotypes such as inflammation, EMT, proliferation, and apoptosis, and rigorously calibrates and validates the model using independent gene expression datasets and machine learning. This comprehensive framework enables robust simulation of disease mechanisms and the identification of combinational therapeutic targets across diverse CRC contexts, extending beyond gene or pathway identification to phenotype-level prediction and systematic intervention analysis.

Additionally, the predictive value of the identified disease signatures was confirmed using supervised machine learning models trained on independent gene expression datasets. The high accuracy and AU-ROC values achieved by these models underscore the robustness of our signatures in distinguishing tumor from healthy tissue, supporting their potential utility as diagnostic biomarkers for CRC. This integrative validation, combining mechanistic modeling, patient survival analysis, and ML classification, demonstrates the translational relevance and reliability of our computational approach.

While the constructed MIM offers a valuable resource for integrating and visualizing CRC-related molecular information, it does not encompass all possible biochemical reactions and pathways involved in CRC. The Boolean modeling framework, while effective for qualitative analysis and hypothesis generation, cannot capture quantitative or graded biological responses. Additionally, our models are specific to the experimental conditions and datasets used for calibration; however, they can be readily adapted to other experimental contexts or datasets as needed. Future work could address these limitations by integrating quantitative modeling approaches, incorporating single-cell and spatial transcriptomics data, and validating predictions in experimental and clinical settings.

In conclusion, our integrative computational approach provides a mechanistic framework for understanding CRC progression and identifying clinically relevant disease signatures and therapeutic targets. The resources generated, including the publicly available MIM and Boolean models, offer a foundation for future mechanistic studies, experimental validation, and the development of personalized combination therapies in colorectal cancer.

## Material and Methods

### Construction of the CRC molecular interaction map

We conducted an intensive literature review to identify critical signaling pathways involved in colorectal cancer (CRC) initiation and progression, including TNF, TGFB, TLR4, WNT, EGF, and Estrogen signaling. Pathway components and their interactions were systematically extracted from peer-reviewed publications (Figure 1), and to capture tumor-immune microenvironment crosstalk, we integrated an inflammatory network from Lu et al. [8]. The map was constructed using CellDesigner [96], adhering to standardized formats: SBML (Systems Biology Markup Language) [97] for computational analysis and SBGN (Systems Biology Graphical Notation) [98] for visual representation. Nodes in the map represent biological entities annotated with official gene symbols (HUGO Gene Nomenclature Committee) and UniProt IDs for protein standardization, while edges define regulatory interactions such as phosphorylation, ubiquitination, activation, or inhibition, and are annotated with PubMed IDs referencing source literature and interaction type. The fully annotated map is publicly available as an interactive web version (https://vcells.net/colorectal-cancer/) and as a downloadable CellDesigner file https://zenodo.org/record/7104858.

### Construction and calibration of the dynamical model

The CRC MIM was converted into Boolean models to perform dynamic systems analysis. The model was divided into three layers: 1) input layer, 2) regulatory layer, and 3) output layer. The input layer and regulatory layer were encoded into Boolean logic. The Boolean model represents a node in one of two possible states: 1 (ON, activation) or 0 (OFF, inactivation). The regulatory relationship from upstream nodes (regulators) to downstream nodes (targets) is expressed by logical operators AND, OR and NOT (Figure 5). NOT gate encodes inhibitory relation. OR gate is used to express the relationship when a target is regulated by multiple regulators independently, i.e., the target will be active if any one of the regulators is active. AND gate encodes the collective effect of multiple regulators on a target. In Boolean models, the future state (t+1) of a node determines by Boolean functions of all its regulators at present state (t): X_i_(t+1) = BF(X_1_(t), X_2_(t),…,X_n_(t)). In Boolean models, the time variable is discrete and usually indicated as a step. To propagate a signal in the Boolean model, different node states update schemes have been proposed, such as synchronous (all nodes updated at each time step) and asynchronous (randomly selected nodes are updated at each time step). Here, we used a synchronous update scheme due to: 1) it does not require detailed biological information for the sequence of events, and 2) it is less computationally demanding, therefore, it can be applied to large biological networks [8,57,58]. We derived Boolean functions of all nodes in the regulatory layer based on their type of relationship (activation or inhibition) with regulators. Further, Boolean functions were calibrated with FC expression data (see Figure 5) and ensure that after steady state (i.e., no change in state variables over time) calculations, nodes in the regulatory layer with negative FCs are represented by 0 states and those with positive FCs are represented by 1 state. For FC expression data, we used two publically available CRC data sets, GSE1323 [37] and GSE8671 [38], from Gene Expression Omnibus (GEO) database. Dataset GSE1323 is a microarray profile of CRC progression to metastasis, containing three SW480 samples (progression) and three SW620 samples (metastatic). GSE8671 is a transcriptome profile of human colorectal adenoma (32 samples) to the mucosa (32 samples) from the same individuals. For each dataset Log2FC expression was calculated using the GEO2R tool (https://www.ncbi.nlm.nih.gov/geo/info/geo2r.html).

The CRC molecular interaction map (MIM) was converted into a Boolean model to enable dynamic systems analysis. The model architecture comprised three layers: (1) input, (2) regulatory, and (3) output. Both the input and regulatory layers were encoded using Boolean logic, where each node represents a biological entity in one of two possible states: 1 (ON/active) or 0 (OFF/inactive) [8,57,58]. Regulatory relationships from upstream regulators to downstream targets were expressed using logical operators AND, OR, and NOT. The NOT gate encodes inhibition, the OR gate represents independent regulation by multiple regulators, and the AND gate captures the collective effect of multiple regulators on a target. Boolean functions for all regulatory layer nodes were derived based on the nature of their regulatory relationships (activation or inhibition). Model calibration was performed using fold change (FC) expression data to ensure that, after reaching steady state, nodes with negative FCs were represented as 0 and those with positive FCs as 1. To illustrate this process, Figure 5 presents a toy network of four nodes (X_1_–X_4_) with both activating and inhibitory interactions. In this example, the network consists of four nodes (X_1_–X_4_) with both activating and inhibitory interactions:

- **Node X_1_** has no upstream regulators and a positive FC value, so it is initialized to state ‘1’ (active).
- **Node X_2_** is regulated by X_1_ (activator) and X_4_ (inhibitor). Although both regulators have positive FC values, X_2_ itself has a negative FC, indicating that both regulators must be active for X_2_ to be repressed; this is encoded with an AND gate.
- **Node X_3_** is regulated by X_2_ and X_4_, both as activators. Here, X_2_ has a negative FC, X_4_ a positive FC, and X_3_ itself is positive. This independent regulatory effect is represented by an OR gate.
- **Node X_4_** is activated solely by X_3_, and both have positive FC values, so X_4_’s future state depends on X_3_’s present state.

After initializing the input node(s), we simulate the network until it reaches a steady state. The steady-state value of each node is then compared to its FC expression value: nodes with positive FCs are expected to have a steady-state value of 1 (active), and those with negative FCs a value of 0 (inactive). This process ensures that the Boolean functions accurately reflect both the regulatory logic and the observed gene expression data.

For calibration, we used two publicly available CRC datasets from the Gene Expression Omnibus (GEO): GSE1323 (microarray profiles of CRC progression and metastasis) [37] and GSE8671 (transcriptome profiles of colorectal adenoma and mucosa from the same individuals) [38]. Log2FC expression values were calculated using the GEO2R tool (https://www.ncbi.nlm.nih.gov/geo/info/geo2r.html).

We encoded phenotypes with multi-valued logic. Inflammation, EMT and apoptosis have five regulators, therefore, they are accepting six ordinal values ranging from 0 to 5, while proliferation has four regulators, thus, it is accepting five ordinal values ranging from 0 to 4. This will allow us to assess quantitatively the effect of multiple regulatory factors on a phenotype. The LSS value of a phenotype is determined by the number of active factors regulating it. For example, if all the regulatory factors are ‘0’, the LSS value of a phenotype is ‘0’. If any one, two, three, or 4 regulators are 1, the LSS value of a phenotype will be ‘1’, ‘2’, ‘3’ or ‘4’ respectively. If all five regulators are ‘1’ simultaneously, the LSS value of a phenotype will be ‘5’.

We encoded phenotypes using multi-valued logic to quantitatively assess the influence of multiple regulatory factors. Specifically, inflammation, EMT, and apoptosis each have five regulators and thus can assume six ordinal values ranging from 0 to 5, while proliferation, regulated by four factors, takes values from 0 to 4. The logical steady-state (LSS) value of a phenotype corresponds to the number of active (state ‘1’) regulators: if all regulators are inactive, the LSS is 0; if one, two, three, or four regulators are active, the LSS is 1, 2, 3, or 4, respectively; and if all five regulators are active, the LSS reaches 5. This approach enables a semi-quantitative assessment of phenotype activation within the Boolean modeling framework [9,99].

### Model simulations and steady-state analysis

Nodes in the input and regulatory layers were encoded using Boolean logic, with each node assigned a state of ‘1’ (active) or ‘0’ (inactive). The initial model state was set by initializing input layer nodes according to the fold-change (FC) expression values of ligand-receptor associations: nodes with positive FC were set to ‘1’, and those with negative FC to ‘0’. Logical steady-state (LSS) analysis was performed using CellNetAnalyzer (CNA) [100], propagating signals from the input layer through the regulatory and output layers until no further state changes occurred. For a given input configuration, the LSS determines the steady-state values of all nodes in the model at the same time step (synchronous update), allowing exploration of network dynamics in response to specific stimulations or perturbations that drive cascades through the system to activate or inhibit phenotypes.

To facilitate interpretation, we calculated aggregate phenotype values by grouping phenotypes into tumor-promoting (inflammation, EMT, and proliferation) and tumor-inhibiting (apoptosis) categories. The aggregate value was computed as:

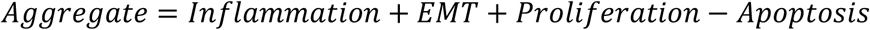

The maximum possible aggregate value is 14, and the minimum is –5.

### Patient data-based validation of disease signatures and drug targets

To validate model-predicted disease signatures and drug targets, we utilized the UCSC Xena browser for survival and expression analyses. Kaplan–Meier survival analysis was performed by stratifying patients based on the expression levels of in silico-derived disease signatures, and calculating overall or progression-free survival probabilities. For survival analysis, patients were grouped into high and low expression cohorts according to signature gene expression thresholds, and statistical significance was assessed using the log-rank test, as implemented in the Xena browser (http://xena.ucsc.edu/).

For validation of the signature involving simultaneous activation of TNF/TNFRSF1A,B and LY96/TLR4 (from GSE8671), we used the TCGA Colon and Rectal Cancer (COADREAD) cohort (https://xenabrowser.net/datapages/?cohort=TCGA%20Colon%20and%20Rectal%20Cancer%20(COADREAD)&removeHub=https%3A%2F%2Fxena.treehouse.gi.ucsc.edu%3A443), dividing patients into high (>13.85) and low (<12.11) expression groups for TNF and TLR4, and calculated progression-free survival. For signatures predicted from GSE1323 (simultaneous activation of TNF/TNFRSF1A,B, EGFR, and inactivation of ERE/ESR), we used the TCGA Colon Cancer (COAD) cohort (https://xenabrowser.net/datapages/?cohort=GDC%20TCGA%20Colon%20Cancer%20(COAD)&removeHub=https%3A%2F%2Fxena.treehouse.gi.ucsc.edu%3A443). Patients were stratified into high and low expression groups for TNFRSF1A, EGFR, and ESR2 (>20.43 and <18.44, respectively), and for TNFRSF1B, EGFR, and ESR2 (>19.45 and <17.54, respectively), with overall survival calculated for each group.

To compare the expression of predicted therapeutic targets between normal and tumor samples, we generated box plots using TCGA (primary colon cancer) and GTEx (normal tissue) data (https://toil-xena-hub.s3.us-east-1.amazonaws.com/latest/GTEX_phenotype.gz; Full metadata). This approach enabled visualization and statistical comparison of gene expression levels in tumor versus normal cohorts, supporting the translational relevance of the identified drug targets. All datasets and visualization tools are publicly accessible through UCSC Xena (http://xena.ucsc.edu/).

### Machin-learning based validation of disease signatures

We retrieved processed gene expression data from 307 healthy tissue samples taken from the GTEX Colon data set (www.gtexportal.org) and 287 tumor tissue samples taken from the TCGA Colon Adenocarcinoma dataset (https://tcga.xenahubs.net) and extracted expression data for the identified markers TNF, TNFRSF1A,TNFRSF1B,EGFR, TLR4 and ESR2. Next, we split the data into a training (75%) and a test (25%) dataset, keeping the original ratio between the cancer and healthy groups. Using the python scikit-learn package, we utilized the data to train and validate instances of several supervised machine-learning (ML) models (k-nearest neighbour, support vector machine, naïve Bayes and random forest). The models’ performances on both training and testing datasets were reported using the area under the receiver operating characteristic curve (AU-ROC). We also computed the confusion matrices.

## Data availability

The CellDesigner file can be downloaded from https://zenodo.org/record/7104858.

The map is available online at https://vcells.net/colorectal-cancer/, where one can easily overlay data to visualize the expression pattern nodes. For illustration, we provided two datasets GSE1323 and GSE8671.

Boolean models are provided at https://zenodo.org/record/7143340#.Yzx2L4RBwuU in a CellNetAnalyzer format.

## Contributions

OW, JV, and FK, developed and conceptualized the idea. The MIM was constructed by FK, JV and MN. Mathematical model, simulations and interpretation of the results were conducted by FK, MN, VR and JV. Model validations were conducted by FK and JV. The manuscript was drafted by OW, JV, FK, AS, and MN. We acknowledge Martin Eberhardt for uploading the map on https://vcells.net/. This work was supported by the Germany Federal Ministry of Education and Research (BMBF) as part of the SASKit project (FKZ01ZX1903B).

## Supporting information

Brief description of each pathway

