## Supplementary material for "Integrative network modeling of colorectal cancer reveals diagnostic signatures and therapeutic targets": Brief description of each pathway

**Brief description of Pathways in the map**

**TNF Pathway:** The TNF pathway has a multi-faceted role in cancer progression. It promotes proliferation, survival, and immune response, and can induce apoptosis [1]. TNF binds to the TNF receptors (TNFRSF1A, TNFRSF1B) on the cell surface. It initiates the NFKB signaling pathway through activation of the TNF/TRADD/RIP1/TRAF complex, which activates TAK1 [1,2]. TAK1 forms a complex with TAB2,3 (MAP3K7/TAB2,3), which phosphorylates IKKα/β. Upon phosphorylation, IKKα/β degrades IKB (NFKBIA, B), which activates and translocates NFKB into the nucleus, where it activates genes involved in cell survival, inflammation, and invasion.

TAK1 also phosphorylates MKKs (MAP2K3,4,5) to transiently activate p38 (MAPK14) and JNK (MAPK8). The TNF pathway supports apoptosis by activating Casp8 through activation of FADD. Cytotoxic T lymphocytes (CTL) activated by Th1 cells express FASL, which also activates Casp8 through FADD [3]. This Casp8-dependent apoptosis can be blocked by c-Flip, which is one of the transcriptional targets of NFKB [3,4].

Further, the TNF pathway triggers the production of reactive oxygen species (ROS) [5] and activation of COX2 (PTGS2) [6,7]. COX2 can also be activated by the SPHK1/S1P pathway, which is an important regulator of sphingolipid metabolism and has been shown to correlate with cell proliferation and survival (regulating NFKB signaling and inhibiting CERAMIDE-dependent apoptosis) [3–5,7]. SPHK1 can also be activated by ERK (MAPK3), which can also be crosstalk between the TNF and EGFR pathway [8]. COX2 also activates Prostaglandin E2 (PGE2/PTGER2), which activates the pro-survival pathways RAS/RAF/MEK/ERK [9] and PI3K/AKT [10]. Further, activated PGE2/PTGER2 can inhibit GSK3β by phosphorylating and promoting β-catenin signaling, which plays an important role in tumor survival and invasion [11].

ROS activates the ATM pathway, which promotes apoptosis by inhibiting MDM2 and activating TP53 [12–14]. ROS is also known to activate the ERK pathway through the activation of MEK1,2 [15]. Further, ROS can activate ASK1 (MAP3K5), which further activates JNK [16]. JNK is a part of AP-1 transcriptional activity, which is involved in the progression of intestinal carcinogenesis [17,18]. In addition, the TNF pathway also activates JNK by activating MEKK1 (MAP3K1) [4].

It is well established that the TNF pathway plays an important role in immune response, and the TNF-mediated intestinal dendritic cell (DC) survival and maturation are presented in the map as activating reactions from TNF/TNFRSF1A, B to the DC node [3,19,20].

**TGF-b pathway:**

Around 80% of colon cancers have a mutation in the TGFB pathway [21]. The TGFB pathway plays a dual role in CRC carcinogenesis. In normal colonic epithelium, TGFB serves as a tumor suppressor by inhibiting proliferation and inducing apoptosis. However, during the later stages of CRC, TGFB acts as a tumor promoter and supports malignant transformation [22]. In the map, the node TFGF1,2,3/TGFBR1,2,3 represents ligand-receptor binding. Upon ligand binding, the TGFB pathway is activated and triggers the activation of SMAD2 and SMAD3, which subsequently bind to SMAD4 to form heteromeric complexes [22,23]. The SMAD2/3/4 complex translocates into the nucleus, where it regulates the gene expression of its targets. The response of the TGFB signaling pathway is highly dependent on the activation and cooperativity of various SMAD molecules [24]. The SMAD2/3/4 complex activates various EMT regulatory elements, including SNAI1,2, ZEB1,2, and TWIST [25]. It can also directly interact with CTNNB1/LEF1 and mediate EMT [25]. SMAD3/4, in combination with FOXO1, activates cyclin-dependent kinase inhibitor 1A (CDKN1A) [26]. SMAD7 inhibits the formation of the SMAD2/3 complex, suggesting its inhibitory role in the TGFB pathway; however, it also promotes TGFB-induced activation of the JNK pathway through non-complex binding SMAD3 [23]. Further, SMAD7 can also interact with β-catenin and may act as a promoter of cell cycle arrest [27].

In addition, TGFB also activates JNK pathways independent of SMADs by activating MAP3K1 and the TRAF6/TAK complex [23,28,29]. The TGFB pathway mediates bone morphogenetic protein (BMP) pathway responses through activation of the SMAD1/5/8 complex [30]. The SMAD1/5/8 complex combines with SMAD4, which subsequently translocates to the nucleus. The SMAD1/5/8/4 complex is inhibited by SMAD6 [30]. BMP is known for its role in promoting growth in colon carcinomas at advanced stages [31,32].

TGFB pathway stimulation also activates the ERK MAPK pathway by phosphorylating ShcA proteins, which form a complex with GRB2 and SOS1 and subsequently can trigger the ERK MAPK pathway [23]. Additionally, TGFB is one of the immunosuppressive cytokines that inhibits TH1 and TH2 cell responses, allowing tumors to escape immune surveillance mediated by CD8+ cytotoxic T lymphocytes [3,33,34].

**TLR4 Pathway:** The TLR signaling pathway is involved in both innate and adaptive immune responses and plays a crucial role in inflammation-associated cancers such as colorectal cancer [35,36]. TLRs recognize a wide array of pathogens and subsequently regulate the expression of key signaling molecules including NF-κB, AP1, STATs, and IRFs, that are involved in immune responses and cell survival [37,38,35]. Dysregulation of this signaling machinery can disrupt epithelial homeostasis and induce chronic inflammation, ultimately leading to colorectal cancer. Among the TLR family members, TLR2 and TLR4 possess a strong ability to recognize a diverse range of pathogenic ligands [39].

In this pathway map, we describe the molecular interaction details of TLR4 signaling. Upon lipopolysaccharide (LPS) binding, the TLR4 pathway is activated, with MD2 (LY96) and MAL acting as accessory proteins to facilitate LPS/TLR4 binding [37]. This activation is mediated by MYD88 and TRAM, which initiate downstream signaling cascades to mount an inflammatory response. The

MYD88-dependent signaling cascade proceeds through IRAKs, TRAF6, and the TAK/TAB complexes, leading to phosphorylation of IKKα/β and MKKs, ultimately resulting in NF-κB activation [37,38,35]. In parallel, TAK/TAB also phosphorylate MKKs (MAP2K3, 4, 6), which activate the JNK/p38 pathways [40].
The TRAM/TRIF complex activates the TLR4 pathway independently of MYD88, initiating a signaling cascade through RIP that converges with TRAF6 and TAK/TAB complexes, resulting in activation of both NF-κB and the JNK/p38 pathways. Additionally, the TLR4/TRAM-TRIF axis can activate IRF3 via TBK1/IKKε to mount an antiviral response. TBK1/IKKε also contributes to NF-κB activation [41].

**Wnt signaling pathways:** Wnt signalling was initially identified for its role in carcinogenesis and was later recognized as essential in embryonic development, including body axis patterning, differentiation, proliferation, and migration [42]. Numerous experimental studies indicate that most colorectal cancers exhibit aberrant activation of the Wnt pathway, frequently due to loss of APC, which is widely considered the initiating and driving event [43].
The Wnt pathway is activated when Wnt ligands bind to receptors of the Frizzled family and LRP receptor complexes [44]. This interaction leads to phosphorylation of Dishevelled (Dvl), disrupting the destruction complex composed of Axin, APC, GSK3B, CHKA, and CTNNB. In the absence of Wnt, this destruction complex mediates ubiquitination and degradation of β-catenin. However, upon pathway activation, the complex is disrupted, allowing β-catenin to accumulate and translocate into the nucleus (for detailed mechanisms, see [45–47]).
Within the nucleus, β-catenin forms a transcriptional complex with TCF/LEF (T-cell factor/lymphoid enhancer factor), driving the expression of target genes such as *CCND1, JUN, MYC, MDM2, VIM, FN1, SNAI, TWIST,* and *ZEB*, while repressing *CDH1*. Wnt signalling also activates the JNK pathway via DVL1 and RAC1, contributing to cell proliferation and migration [48]. In addition, the inhibitory effect of Wnt-5a on canonical Wnt signalling has been incorporated into the map, as it is proposed to suppress β-catenin activity [48,49].

**EGFR signaling pathways:** Mutations that induce aberrant activation of the epidermal growth factor receptor (EGFR) pathway have been associated with many cancer types and are already targeted by several anticancer therapeutics [50]. EGFR receptors and their downstream pathways are important regulators of colorectal cancer development and progression [51]. Upon ligand binding, the EGFR receptor initiates several signal transduction cascades, including the mitogen-activated protein kinase (MAPK) and the phosphatidylinositol 3-kinase (PI3K)-AKT pathways, modulating cell migration, proliferation, evasion of apoptosis, and immune responses [51,52].
EGFR activates the Ras/RAF/MEK/ERK signaling pathway, which subsequently activates MYC, RSK, MSK, SPHK1, and STAT3, proteins shown to correlate with cell proliferation and survival [53,54]. The EGFR-dependent activation of the PI3K/AKT and JAK/STAT pathways results in regulatory factors

involved in survival, anti-apoptosis, and migration. For example, the PI3K/AKT pathway activates the protein synthesis factor mTOR/S6K and anti-apoptotic factors such as BCL2 [55]. The PI3K/AKT pathway also protects cells from apoptosis by activating the anti-apoptotic factor XIAP through NFKB1, and further inhibits apoptosis by suppressing p73 and activating MDM2 to degrade p53 [3].

In addition, the JAK/STAT pathway, via STAT3, promotes the transcription of genes such as MYC, CCND1, BCL2L1, SOD (which inhibits ROS), IL6, SNAIs, VIM, and MMPs, thus regulating proliferation, immune response, anti-apoptosis, and migration [56,3]. Furthermore, the EGFR pathway also regulates autophagy (the natural mechanism for clearing out damaged cells) by disrupting the Beclin 1/BCL2 complex to release Beclin 1 and induce autophagy [3].

**Estrogen signaling pathways:** Men are more likely to develop colorectal cancer compared to women of similar age, suggesting a protective effect of sex hormones in the development of the disease [57,58]. The estrogen pathway is activated by binding estrogens to specific estrogen receptors (ERs), which stimulate transcription or initiate signaling cascades that activate or inhibit the expression of genes. In this map, we included the molecular events governing the regulation of genes by estrogen receptor alpha (ERα) and estrogen receptor beta (ERβ). ERα and ERβ have distinctive effects in CRC, with ERα acting as a tumor promoter while ERβ protects against tumor development [59,60]. Further, the biological response of cells to estrogens depends on the ratio of ERα to ERβ, and ERβ inhibits the transcriptional activity of ERα [57,61]. Advanced-stage CRC is associated with the loss of ERβ [62], and therapies have been designed to treat CRC using ERβ agonists [63].

Upon estrogen binding to the ERα receptor, the PI3K/AKT pathway is activated, which leads to phosphorylation of ERα and its translocation into the nucleus to regulate the transcription of genes including SP1, JUN, FOS, and NFKB, key factors in proliferation and cell survival [57]. ERα can also be phosphorylated without estrogen ligand binding, through the receptor tyrosine kinase (RTK) pathway (e.g., EGFR) via Ras/Raf/ERK1,2 and RSK (ribosomal protein S6 kinase) signaling [57]. In contrast, ERβ-mediated signaling inhibits proliferation and survival by activating diverse intracellular pathways, including protein kinase C (PKC) and Ca2+ pathways [57]. ERβ induces Ca2+ influx into the cell and suppresses PKC signaling, thereby inhibiting PI3K/AKT-mediated cell proliferation and survival [57]. ERβ-mediated PKC-Ca2+ signaling can also inhibit FOXC2 expression, previously identified as an important regulator of breast cancer migration and invasion by activating β-catenin and ZEB [64,65]. Additionally, ERβ can modulate the cell cycle by inhibiting CCND1 and MYC and activating p21 and p53, thereby suppressing cell cycle progression [59,66]. Further, ERβ induces an anti-inflammatory response in colorectal cells by inhibiting the IL6 pathway [59].

**Immune microenvironment network:** Advancements in biomedical research and high-throughput technologies have revealed that cancer and inflammation are closely correlated [67,68]. In the last few decades, the role of the immune system and inflammation in cancer development and progression has gained enormous research interest. The aberrant immune response in inflammatory bowel disease (IBD), which progresses to CRC in many patients [69], involves hyperactivation of innate and adaptive immunity [69]. Emerging evidence suggests that dysregulation of the immune microenvironment and inflammation-related pathways—including TNF, TGFB, TLRs, IL6, STAT3, COX2/PGE2—has contributed to the development of inflammation-associated cancers, such as colorectal cancer [3,70–74]. To study the link between immune system inflammation and colorectal cancer, we included an immune microenvironment network (highly interconnected and containing many feedback loops) from Lu et al. [3] in this CRC map.

The immune response can be initiated by antigen-sensing cells such as dendritic cells (DC) [75], which are activated by cytokines and chemokines such as tumor necrosis factor (TNF) and CCL2, respectively [19,20]. DCs activate IL-12 to trigger the Th1 immune response, which subsequently mediates the activation of CD8+ cytotoxic T cells (CTL) through IFNG, a critical factor in immune surveillance [76]. Th1 cells also mediate the activation of macrophages (MAC) via IFNG, leading to production of pro-inflammatory cytokines such as TNF-α (TNFA) and IL-6 [77,78]. IL-6 can also be activated by DCs in IBD progression [79]. Furthermore, DCs can secrete IL-4 to initiate humoral immunity by activating Th2 cells (TH2) [80,81].

Th1 and Th2 cell responses are counteractive: Th2 inhibits the Th1 response via cytokines such as IL-4 and IL-10, while Th1 inhibits the Th2 response using cytokines such as IFNG [82]. Additionally, DCs mediate the differentiation of regulatory T cells (TREG), triggering the activation of immune suppressive cytokines including TGFβ and IL-10, which have been shown to lessen IBD [3,74].

Genes responsible for pro-inflammatory cytokines such as IL-6 and TNFA can also be activated by the JNK/p38 [83] and NFKB [84] pathways. The IL-6–mediated JAK/STAT pathway in CRC is involved in enhancing cell proliferation and tumor growth [85].

**Interconnectivity among pathways:** Pathways in a cell are interconnected rather than independent and can affect each other considerably through crosstalk. The role of crosstalk among pathways can be a double-edged sword: it can ensure homeostasis against perturbations, but on the other hand, it can also make these pathways susceptible to gain-of-function mutations and constitutive signal activations, which interferes with drug efficacy due to redundant links that negate the drug effect [86]. The interconnectivity among pathways poses a challenge to single-drug therapy, which may be counteracted by combination therapy. By integrating various pathways in this CRC map, we observed that these pathways interconnect and crosstalk at multiple points and levels. For example, the STAT3 oncogenic signaling pathway establishes several levels of crosstalk (e.g., STAT3-PTGS2, IL6-STAT3) between tumour cells and their immune microenvironment [87].

PTGS2 interacts with EGF and downstream signaling pathways by mediating the activation of KRAS/RAF/MEK/ER and PI3K/AKT signaling [3,10,88]. It has been reported that PTGS2 and EGF synergistically promote CRC progression and metastasis [89]. Additionally, PTGS2 also interacts with the WNT pathway by promoting β-catenin signaling [3,11]. PI3K/AKT signaling is also mediated by TNF signaling [90]. Another observed crosstalk in the map is the TGF signaling pathway with MAPK and WNT pathways through SMAD2/3, ERK, JNK, P38, and β-catenin signaling to promote CRC progression [23].
